# Temporal patterns in electrical nerve stimulation: burst gap code shapes tactile frequency perception

**DOI:** 10.1101/2020.04.10.033241

**Authors:** Kevin K. W. Ng, Christoffer Olausson, Richard M. Vickery, Ingvars Birznieks

**Affiliations:** School of Medical Sciences, UNSW Sydney, Sydney, Australia; Neuroscience Research Australia, Sydney, Australia; Faculty of Medicine and Health Sciences, Linköping University, Linköping, Sweden

## Abstract

We have previously described a novel temporal encoding mechanism in the somatosensory system, where mechanical pulses grouped into periodic bursts create a perceived tactile frequency based on the duration of the silent gap between bursts, rather than the mean rate or the periodicity. This coding strategy may offer new opportunities for transmitting information to the brain using various sensory neural prostheses and haptic interfaces. However, it was not known whether the same coding mechanisms apply when using electrical stimulation, which recruits a different spectrum of afferents. Here, we demonstrate that the predictions of the burst gap coding model for frequency perception apply to burst stimuli delivered with electrical pulses, re-emphasising the importance of the temporal structure of spike patterns in neural processing and perception of tactile stimuli. Reciprocally, the electrical stimulation data confirm that the results observed with mechanical stimulation do indeed depend on neural processing mechanisms in the central nervous system, and are not due to skin mechanical factors and resulting patterns of afferent activation.

## Introduction

Temporal features of neural spike activity play a major role in encoding tactile information [1–4]. In a previous study, we used pulsatile mechanical stimulation to investigate the neural basis of frequency perception in the flutter range. When we grouped pulsatile stimulation patterns into periodic bursts, we found that perceived frequency was determined by the duration of the silent gap between bursts of spikes in peripheral afferents (termed the burst gap code), irrespective of the number of spikes within a burst or the burst rate [5]. This stands in contrast to the mean spike rate or periodicity code that are often proposed to be neural codes for frequency perception in higher neural centres [6–9]. Electrical burst stimulation has been suggested to be useful in restoring the sense of touch in amputees using a prosthesis [10–12] or in the treatment of movement disorders through deep brain stimulation [13–15]. These studies were generally concerned with the efficacy of inducing neural activity, rather than the effect of burst stimulation on information encoding. We believe that careful consideration of neural coding related to the temporal aspects of these burst patterns could potentially improve the performance of such stimulation strategies.

The discovery of the burst gap code for perceived frequency depended on the use of brief mechanical pulses that each only evoke a single spike in responding afferents, as had been verified using microneurography, allowing us to precisely control the spiking pattern of responding tactile afferents [5, 16, 17]. Electrical stimulation activates peripheral axons in a manner different than that of mechanical stimulation [18], by bypassing the specialised mechanoreceptors in the skin and directly stimulating the tactile afferent axons [19, 20]. Electrocutaneous stimulation non-selectively activates the different types of tactile afferents (associated with different skin receptors) [21, 22], which contrasts with mechanical stimulation where the type of afferent recruited varies depending on the sensitivity of the fibre type to stimulus characteristics such as the vibration frequency of a mechanical sinusoid [23, 24].

A single tap on a finger generates a complex propagating wave on the fingertip skin [25, 26] that will evoke complex and potentially variable population spike patterns due to sporadically responding afferents [27]. In particular, such a stimulus would likely reliably activate afferents whose receptive fields are directly under the probe, but this activation would become stochastic at some distance from the locus of stimulation [24, 28]. This raises the possibility that the burst gap code that we observed with mechanical stimulation is in some way dependent on these spatial activation patterns and/or predominant activation of certain afferent types. As an alternative, electrical stimulation is a robust method to reliably activate axons, and a simple technique to implement in neuroprosthetic interfaces [29] or haptic displays [30]. Thus, in this study we aimed to use electrical stimuli to directly evoke spike trains in tactile afferent fibres while avoiding complex mechanical skin phenomena and establish the burst gap model for frequency perception.

## Materials and methods

### Subjects

Two separate experiments were conducted. Twelve subjects (aged 21-26, 5 females) participated in Experiment 1. Eight subjects (aged 21-25, 5 females) participated in Experiment 2. Three of the subjects participated in both experiments. All subjects were healthy and without any known history of altered tactile function. The experimental protocols were approved by the Human Research Ethics Committee (HC16245) of UNSW Sydney and written consent was obtained from subjects before experimentation began.

### Apparatus

Stimulus patterns were generated via a CED Power1401 mk II data acquisition system using Spike2 (Cambridge Electronic Design, Cambridge, UK) and MATLAB (Mathworks, Natick, USA) software. These patterns triggered constant current electrical pulses from a DS7AH stimulator (Digitimer, Welwyn Garden City, UK). The electrical stimuli were delivered to the right index finger of subjects using Kendall 200 Series foam electrodes (Covidien, Mansfield, MA, USA), with one electrode placed on the distal interphalangeal joint and another on the proximal phalanx, to stimulate the digital nerve. A button box was used by subjects to indicate their responses to the psychophysical task. Compound action potentials were recorded using a PowerLab data acquisition system and LabChart software (ADInstruments, Sydney, Australia).

### Stimulation patterns

#### Experiment 1

The stimulus timing patterns used were based on that of our previous study [5]. In our first experiment, the aim was to determine whether burst gap coding would best describe the perception of frequency in subjects. The test stimuli consisted of one of four trains with periodic bursts of 2-4 pulses (Fig 1A), such as would be generated in the afferent response to high-amplitude vibration [24, 31]. The individual pulses within the burst were spaced either 4.35 ms (stimuli a-c) or 8.7 ms (stimulus d) apart. The period between repeating bursts was fixed at 43.5 ms, resulting in a burst rate of 23 Hz. However, the mean pulse rate and the inter-burst interval (t_L)_ varied noticeably between the four stimuli, ranging from 46 to 92 pulses/s and 26.1-39.15 ms, respectively.

**Fig 1.**
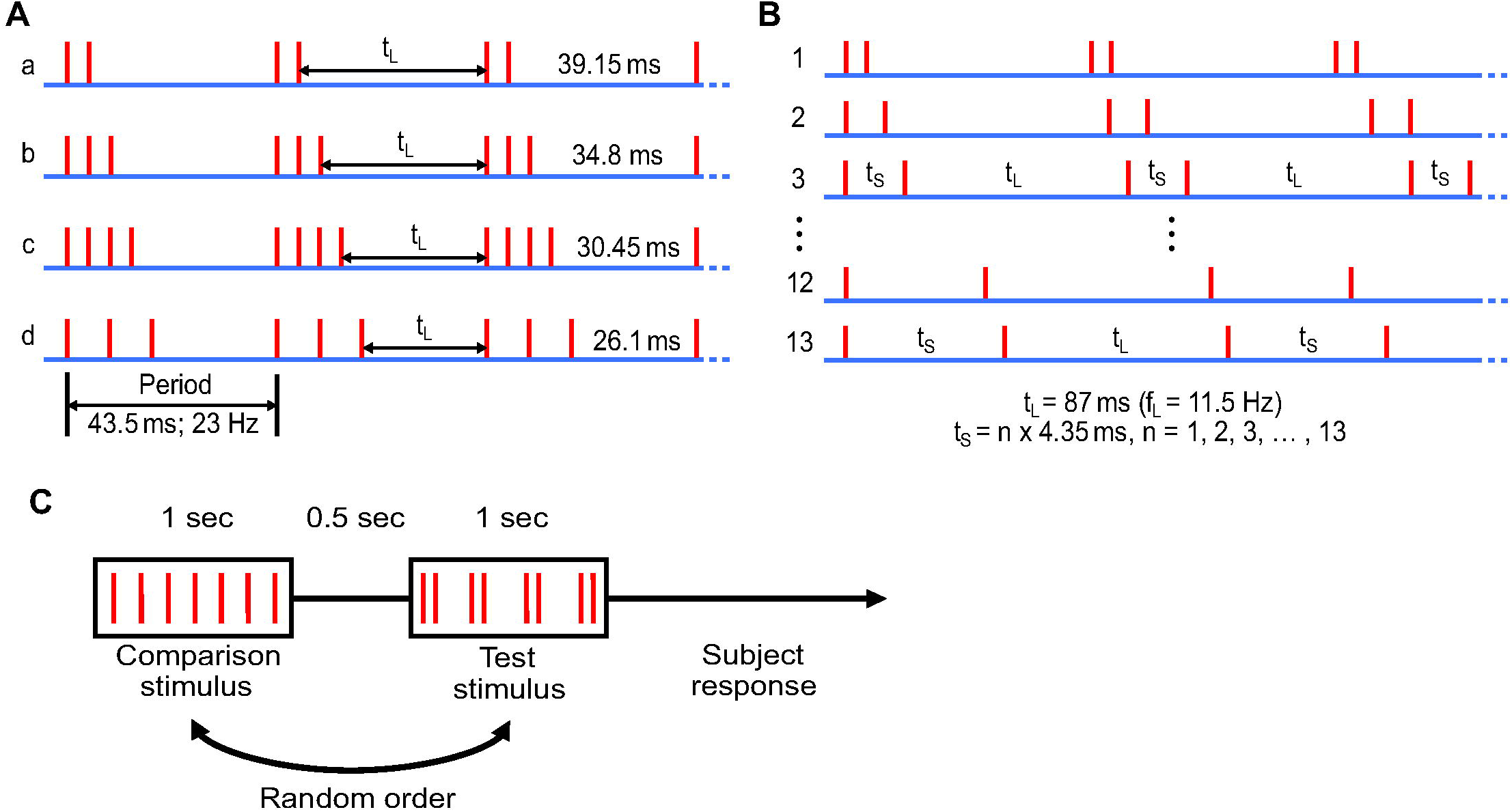
Schematic of stimulus patterns and experimental protocol, each red vertical line represents an electrical pulse. (A) In Experiment 1, test stimuli were pulse trains with periodic bursts of 2-4 pulses. The stimuli have the same periodicity of 43.5 ms (burst rate of 23 Hz), but different inter-burst intervals (t_L)_ and overall pulse rates. (B) In Experiment 2, test stimuli consisted of a short interval doublet (t_S)_, followed by an 87 ms long inter-pulse interval (t_L)_. The short inter-pulse interval was varied in steps of 4.35 ms, from 4.35 ms up to 56.55 ms. (C) Two interval forced choice procedure used to determine perceived frequency of test stimuli.

#### Experiment 2

In our second experiment, we measured the critical value of the inter-pulse interval within a burst that determined how the pulses contribute to perceived frequency. The test stimuli were one of 13 trains of a 2-pulse burst (doublet) with a short interval (t_S_) and a long interval (t_L)_ (Fig 1B). The short inter-pulse interval (t_S)_ ranged from 4.35 ms to 56.55 ms, increasing in steps of 4.35 ms. The long interval (t_L)_ was fixed at 87 ms. The burst gap code predicts that when two pulses are close enough in time, they will be treated as a burst and accordingly the perceived frequency will be determined by the burst gap (approximately 11.5 Hz in this experiment).

### Electrical stimulation parameters

Electrical stimuli were 0.1 ms wide square pulses. Prior to the start of each experiment, the electrical current level was optimised for the subject so that the electrical pulses were perceived clearly as a series of taps without discomfort. Current levels used ranged from 6.5 to 11.5 mA across subjects and were kept at a fixed level within a session.

### Verifying the stable activation of a population of afferents

To verify that the afferent axons fired reliably to each electrical pulse within a burst in our closely spaced stimulation patterns, sensory nerve action potentials (SNAPs) were measured in two subjects by stimulating the sensory nerves distally (as described above) and recording the compound action potential more proximally [32]. The recording electrodes were located just proximal to the wrist crease over the median nerve, with the ground electrode placed on the thenar eminence. SNAPs were recorded from 200 repetitions of the burst patterns used in Experiment 1 (Fig 1A).

### Measuring the point of subjective equality for frequency

To determine the perceived frequency of each test stimulus, a two-interval forced choice paradigm was used as in our previous study [5]. Test stimuli were compared against 6-8 different comparison stimuli, which were regular trains of pulses with repetition rates ranging from 14-54 Hz for Experiment 1 and 7-21 Hz for Experiment 2. For each trial, a test stimulus and a comparison stimulus were presented for 1 s each, in a random order and separated by 500 ms (Fig 1C). The subject indicated by button press which of the two stimuli had the higher frequency. Following optimisation of the electrical current, a practice round was conducted which comprised a selection of test and comparison stimuli to familiarise subjects with the task, before the actual test procedure began. To obtain psychometric curves, each test condition was compared 20 times against each of the regular comparison stimuli, giving rise to 120-160 trials per test stimulus.

At each comparison frequency, we determined the proportion of times the subject indicated that the comparison stimulus had a higher frequency than the test stimulus (P_H)_. To obtain a linear psychometric function, the logit transformation ln(P_H_/(1−P_H_)) was applied to the data [33]. The value of the comparison frequency that is equally likely to be judged as higher or lower than the test stimulus was taken as the point of subjective equality (PSE). This value corresponds to the frequency where the zero crossing of the logit axis occurs for the regression line fitted to the logit transformed data (example in S1 Fig).

### Statistical analysis

For Experiment 1, linear regression was performed to determine if the slope of the line fitted to the data would be better explained by the burst gap (i.e. the inter-burst interval) or the burst rate (periodicity), which was constant at 23 Hz. A two-way ANOVA (repeated measures within the same subject for the different stimulation methods) was used to determine if there were differences between the data collected here using electrical stimulation and the mechanical data collected in our previous study. For Experiment 2, a joinpoint regression model created using Joinpoint Trend Analysis Software (National Cancer Institute, Bethesda, MD, USA) was used to determine the trend that best fitted the psychophysical data. An alpha level of 0.05 was used for all statistical tests.

## Results

### Stable activation of the afferent population

Electrical stimulation with patterns used in Experiment 1 (Fig 1A) resulted in SNAPs that were consistent within a burst irrespective of inter-pulse intervals used or the number of pulses within a burst (Fig 2, Fig 3A). To examine pulse-to-pulse variability within individual bursts, we normalised the amplitude of subsequent SNAPs to that of the first SNAP for each burst; this showed that the nerve response to pulses occurring later in a burst were not negatively affected by preceding pulses (Fig 3B), which suggests that the stimulus burst patterns were faithfully reproduced in the nerve firing patterns.

**Fig 2.**
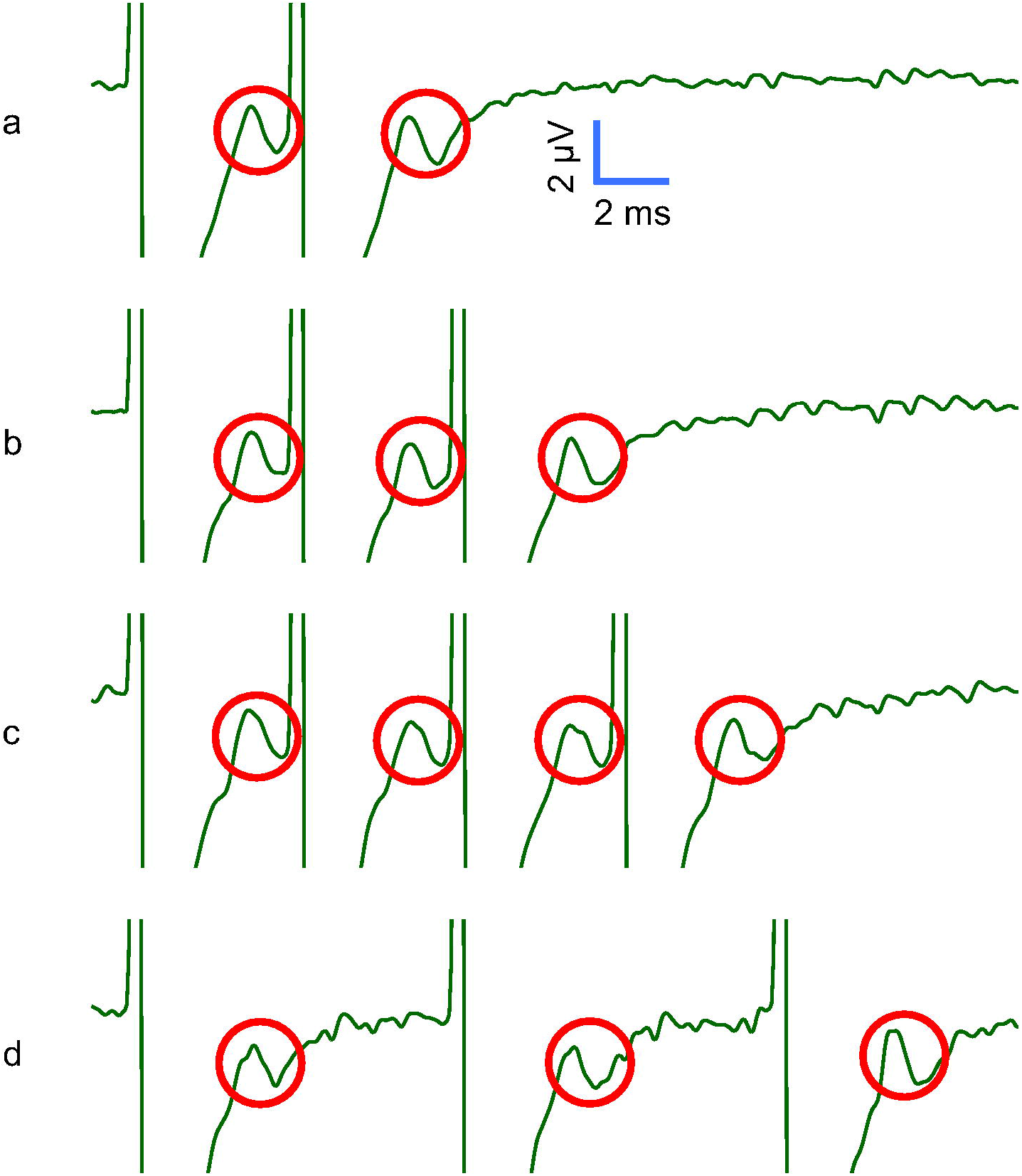
Sensory nerve action potential (SNAP) recordings. Average of 200 compound action potentials recorded from the median nerve in one subject, in response to Experiment 1 stimulation patterns illustrated in Fig 1A. The red circles indicate SNAPs, while the clipped waveform occurring before each SNAP represents the electrical stimulation artefact.

**Fig 3.**
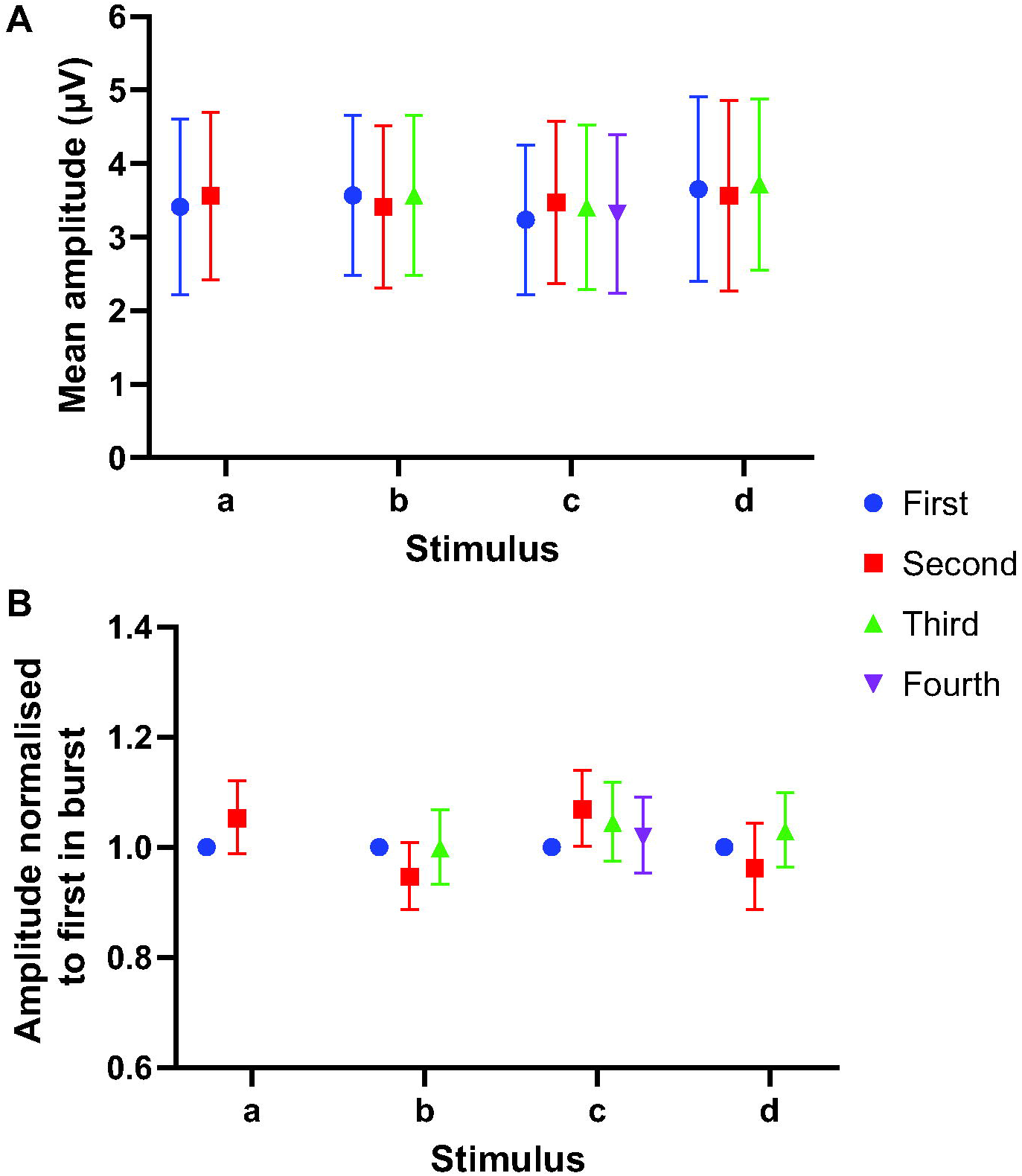
SNAP amplitudes within a burst. (A) The mean peak-to-peak amplitude (n = 200) of evoked SNAPs in one subject. Error bars denote standard deviation. (B) Geometric mean of SNAP amplitudes after normalising to the first SNAP within each burst response. Error bars denote 95% confidence intervals.

### How is perceived frequency encoded?

In Experiment 1, the aim was to determine which coding mechanism could best describe the perception of frequency in subjects when the electrical pulses were grouped into bursts. The mean data from the 12 subjects are shown in Fig 4. The observed perceived frequencies were 2- to 4-fold lower than that predicted by the mean pulse rate (Fig 4A green triangles), indicating that a simple rate code could not explain perceived frequency in subjects. Linear regression was performed between the reciprocal of the burst gap (t_L)_ and the perceived frequency for each stimulus (Fig 4B). The regression line had a slope that was non-zero (1.2, 95% CI 1.0-1.4; R^2^ = 0.76; p < 0.0001), which does not correspond to that of the burst rate / periodicity (expected slope 0, Fig. 4A pink triangles), but is a good match to the reciprocal of the burst gap (t_L)_. The slope here closely resembles that of the mechanical data from our previous study (slope = 1.3, 95% CI 1.0-1.5; R^2^ = 0.68; p < 0.0001), which also best matched 1/t_L_ [5].

**Fig 4.**
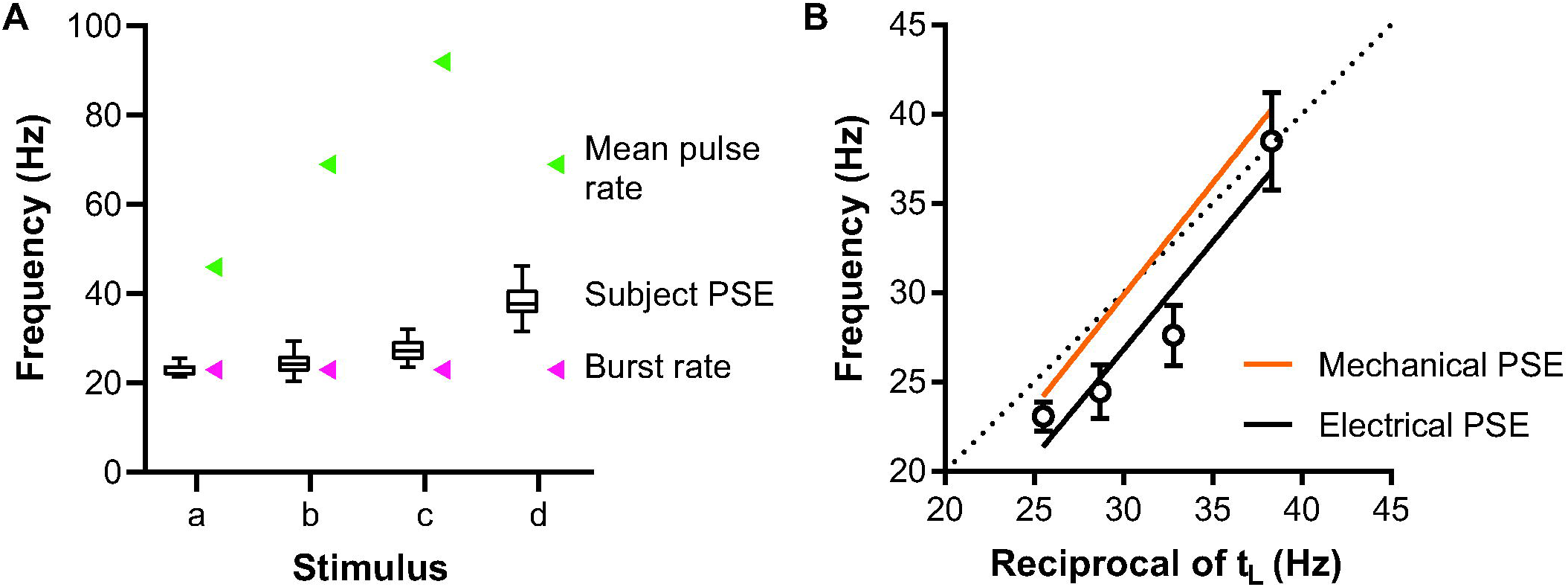
Possible coding schemes for frequency perception of electrical burst stimuli. (A) Boxplots with whiskers represent the subjective perceived frequency of the electrical stimuli used in Experiment 1 (n = 12). Green and pink arrowheads indicate predicted PSE values if the frequency judgement had been based on the mean pulse rate and the burst rate (periodicity), respectively. (B) Fitted regression line (black line) indicating that the mean electrical PSE data (black circles; n = 12) for perceived frequency is a close match to the reciprocal of the burst gap t_L_. The regression line fitted to mechanical data from our previous study [5] is shown for comparison (orange line). The dotted line is the line of identity, where the PSE would equal the reciprocal of the burst gap t_L_. Error bars denote 95% confidence intervals.

A two-way ANOVA indicated that there was a statistically significant main effect by the stimulus pattern (p < 0.0001, F_2,44_ = 143, 71% of total variation) and the method of stimulation (p = 0.0084, F_1,22_ = 8.4, 4.7% of total variation). However, there was no interaction effect (p = 0.36, F_3,66_ = 1.1), suggesting that varying the stimulation pattern had the same effect on frequency perception regardless of whether mechanical or electrical stimulation was used.

### What defines an inter-pulse interval as part of a burst?

For Experiment 2, we varied the duration of the short interval to determine which inter-pulse intervals (t_S)_ would be treated as a burst, so that the perceived frequency depended only on the longer interval (t_L)_. The results from the 8 subjects are shown in Fig 5 which plots the apparent frequency of each stimulus. A joinpoint regression model was fitted, which indicated that the data was best explained by three linear segments. The first segment extended from 4.35 to 13.05 ms, with a slope that was not significantly different from zero (p = 0.64). The next two segments had slopes that were significantly non-zero. The second segment spanned 13.05 to 34.8 ms and had a slope of 0.2 (p = 0.0002). The last segment had a range between 34.8 and 56.55 ms, with a slope of −0.08 (p = 0.0002).

**Fig 5.**
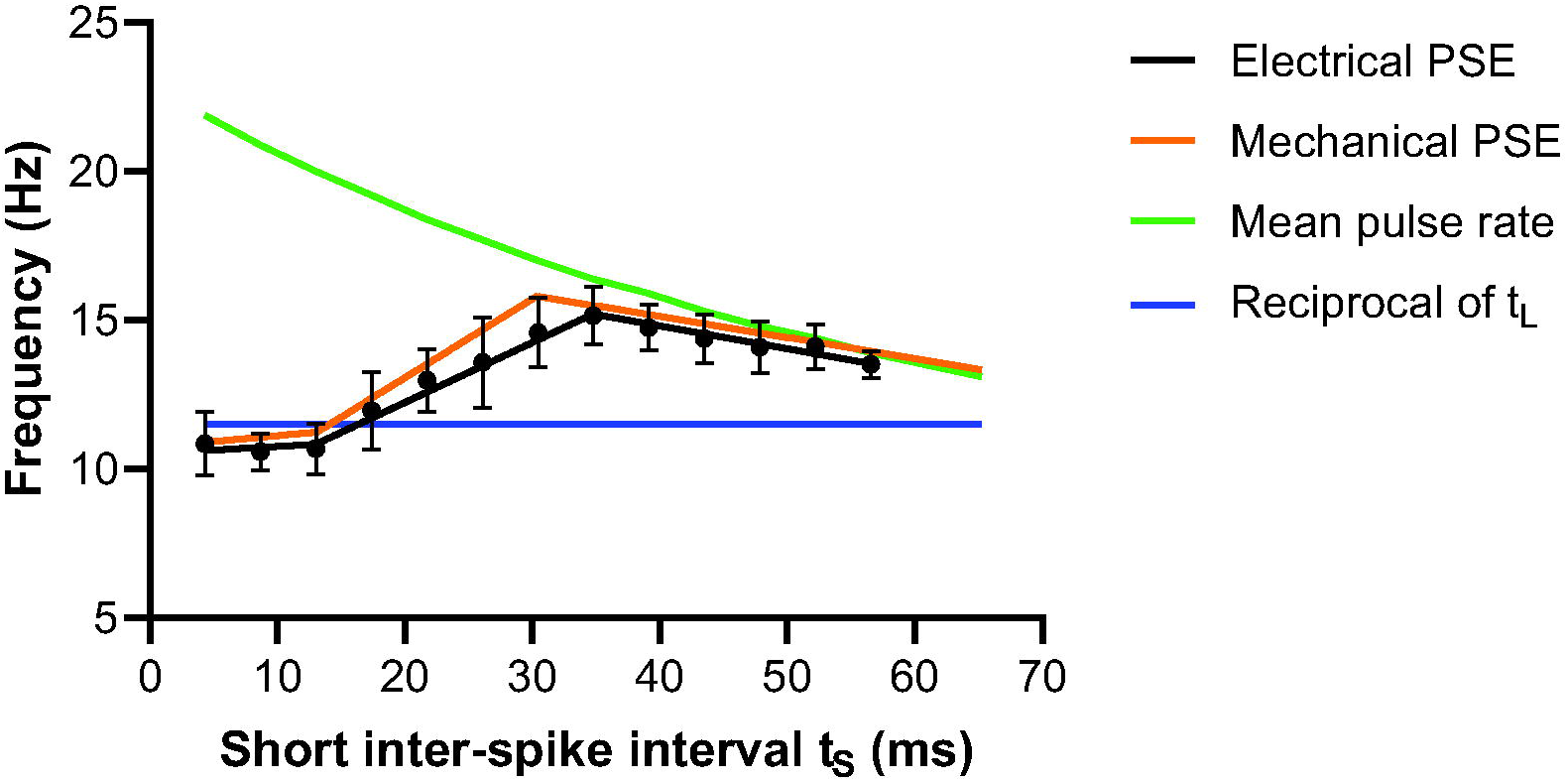
Relationship between the short inter-pulse interval and perceived frequency. PSE values obtained using Experiment 2 stimuli, indicated by black circles (Electrical PSE; n = 8). The black line signifies the three fitted linear segments as determined by the joinpoint regression model. The green and blue lines represent predicted PSE values if frequency judgement had been based on the mean pulse rate and the burst gap code, respectively. The orange line (Mechanical PSE) is the joinpoint model based on the mechanical data from our previous study [5]. Error bars denote 95% confidence intervals.

The slope for the segments here were similar to those reported in our previous study involving mechanical stimulation (Fig 5, Mechanical PSE), though the joinpoint between the second and third segments occurred at 30.45 ms in that study [5], thus at a slightly shorter inter-pulse interval than in the current study (34.8 ms). Nevertheless, the same basic findings were observed in both studies, as shown by the confidence intervals overlapping with the joinpoint model based on the mechanical data and an identical first jointpoint. This suggests that the same definition of a burst can be applied to both mechanical and electrical stimulation.

## Discussion

The main objective of the present study was to verify whether the burst gap coding model for frequency perception could be observed with electrical stimulation in the flutter range. This is an important step to demonstrate due to the potential value of this coding strategy for implementation in bionic systems for providing somatosensory information. Our results show that the burst gap model, based on the duration of the silent inter-burst interval, could reliably predict perceived frequency of complex bursting spike patterns when the spikes were elicited by electrical pulse stimulation of axons. This means that the same overall spike rate in an axon can produce a different frequency percept, depending on how those spikes are arranged in time. This reinforces the importance of the temporal structure of spiking patterns in neural processing and sensory perception.

### Encoding perceived frequency

Electrical stimulation bypasses the complex viscoelastic properties of the skin that may have affected previous work using mechanical stimulation, and so provides strong independent confirmation of the burst gap code. We verified that the efficiency of the electrical stimulus was not affected by the repetition rate by measuring SNAP recordings, which showed that even the shortest inter-pulse interval (4.35 ms) resulted in reliable and consistent activation of tactile afferents. As we used a reasonably strong stimulation current at a supra-threshold level, it was expected that a majority, if not all, of the large myelinated afferents of the digital nerve would be reliably activated regardless of the inter-pulse interval chosen. The ability for the peripheral sensory nerve to faithfully relay the presented burst stimulation patterns, as evident in the SNAP recordings, suggests that perceived frequency depends on the processing of these signals within the central nervous system and not peripheral factors.

While the results in our first experiment, where we used electrical pulses to create bursting trains of the same periodicity but different number of spikes, were best explained by the burst gap code rather than a periodicity or rate code, the mean PSEs were lower than that would be predicted simply from the inter-burst duration. Though the difference in PSEs between electrical and mechanical stimulation was statistically significant, it is unknown whether this difference indicates any physiological importance, as it is well within the Weber fraction of ~0.2-0.3 that has been previously reported in the literature [16, 34–38]. One possible explanation for this bias towards lower frequencies may be that electrical stimulation recruits all afferent types non-selectively, so that the slowly adapting (SA) afferents would be recruited in addition to the fast adapting (FA) afferents, whereas the mechanical stimulus predominantly activated FA afferents [5]. Although previous research has found that SA afferents and their corresponding cortical sensory neurons do not contribute to vibrotactile frequency perception [39–41], their activation and the corresponding sense of pressure they convey [23] may bias subject judgements in psychophysics tests.

These findings are consistent with recent findings from our laboratory which revealed that the spiking pattern, rather than activated afferent type, determined the perceived frequency of repetitive mechanical stimuli [16]. In that study, we found that low-frequency spike trains in Pacinian (FA2) afferents could induce a vibratory precept, even though Meissner (FA1) afferents had been traditionally thought to be solely responsible for perception of low frequency flutter vibrations (<60 Hz), whereas higher frequency activation of Pacinian (FA2) afferents would evoke a different percept of vibratory hum [24]. A generalised frequency processing system that operates with inputs from a variety of receptor type inputs would explain these findings, as well as the constancy of vibrotactile frequency perception across different skin regions innervated by different afferent types [34, 42], e.g. hairy skin lacks Meissner corpuscles [43, 44]. The congruence of the present results using electrical stimulation whereby all types of afferents are recruited non-selectively, with the previous results using mechanical stimuli that activated predominantly FA afferents, supports the suggestion of a generalised frequency processing mechanism.

Natural stimuli encountered during object manipulation and surface exploration generally evoke complex discharge patterns in tactile afferents [2, 45–47]. Additionally, numerous studies have implicated the importance of frequency content in skin vibrations for encoding tactile information such as texture [45, 48, 49]. Accordingly, the frequency perception mechanism substantiated here may represent a more universal method of encoding complex naturalistic patterns of vibration, which may not necessarily have a simple fixed periodicity. For instance, we previously showed that this sensory coding strategy also applied to stimuli that are irregular or aperiodic in nature, as it relies on the processing of individual inter-pulse intervals by weighting their perceptual contribution to perceived frequency depending on their duration, rather than detection of periodicity or spike counting [50].

### The value of bursts

The second experiment, with doublets of different inter-pulse durations, clearly indicate that when the inter-pulse interval for the doublet is sufficiently brief (<15 ms), the pulses are interpreted as belonging to a single burst event and the burst gap determines the perceived frequency. As the separation of the two pulses is increased beyond 15 ms, the propensity for two consecutive pulses to be regarded as a burst decreases and the within-doublet interval appears to have a partial weighted contribution to frequency perception [50]. Beyond 30-35 ms, each individual pulse is treated as a discrete event and the perceived frequency approaches the mean pulse rate.

Burst firing is a neural feature common in various sensory systems [51]. It has been proposed that bursts play a major role in the reliable signalling of neural information, due to its ability to induce long-term synaptic plasticity and to encode more information as compared to single spikes [52, 53]. Moreover, bursting inputs may be required to cause the firing of postsynaptic neurons through interactions at the preferred oscillation or resonance frequency of the particular synapse and may represent a way to increase transmission security of weak inputs [54]. The burst duration reported in such studies ranges between 10-25 ms, which agrees with the 15 ms value reported in our current study.

Burst stimulation has been used as a strategy for delivering sensory information in studies investigating tactile displays [55, 56] and neural prosthetics [10-12]. Using stimulus bursts may provide additional parameters that can be varied in controlling the perception of tactile features such as intensity sensation. Previous studies have varied the width of the electrical pulse, which varies the charge delivered and changes the number of afferents recruited, in turn modulating perceived intensity [37]. However, altering the charge delivered by a pulse risks generating discomfort in the subject, so a method which can extend the range of intensity sensation without changing individual electrical impulses may be of value [57]. Whilst modulating the pulse rate at a fixed pulse width can influence perceived intensity [37], it would also change perceived frequency [57]. The benefit of the burst stimulation method we describe here is that we can control perceived frequency, which is determined by the burst gap, regardless of burst features such as the number of pulses within a burst [5]. Hence, in haptic devices and brain-machine interfaces, this coding strategy may provide a way to modulate frequency perception by means of changing the inter-burst interval and possibly other perceptual features of tactile stimuli encoded by spiking activity within individual bursts [58].

The present study used external electrodes to stimulate the digital nerve and confirm the burst gap phenomenon for frequency encoding, albeit with a small frequency shift compared with mechanical activation. The use of intrafascicular stimulation with multi-electrode arrays could activate nerve fascicles which may have a more restricted range of afferent types, in addition to offering improved spatial resolution [29, 59]. It would be interesting to see whether predominantly FA-containing fascicles show frequency responses that are a closer match to the results with mechanical stimulation in our previous study.

## Conclusions

When stimulus pulses are grouped into periodic bursts, perceived frequency is best explained by the inter-burst interval, i.e. the duration of the silent gap between bursts, as opposed to the burst rate (periodicity) or mean pulse rate. This finding, consistent with what we have shown with mechanical pulsatile stimulation, suggests that this frequency encoding arises from central processing rather than factors such as skin mechanics. The method of burst stimulation described here may represent a way of providing sensory feedback information such as frequency for tactile displays and sensory prostheses.

## Supporting information

**S1 Fig. Example psychometric function.** An example of a logistic regression fit applied to psychometric data from a single subject in one test condition (Experiment 1, stimulus d). PSE (point of subjective equality) is determined as the frequency that gives logit = 0 from the fitted regression line. In this case, PSE was estimated to be 38.1 Hz.

**S2 Dataset. Individual subject PSE values for each experiment.**

