## Supplementary figures and images for "Temporal patterns in electrical nerve stimulation: burst gap code shapes tactile frequency perception"

### S1 Figure

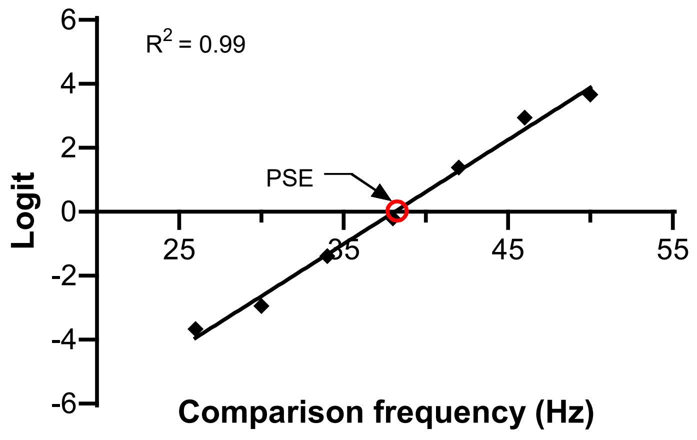
